# Epigenetic variation can promote adaptation by smoothing rugged fitness landscapes

**DOI:** 10.1101/2025.06.06.658353

**Authors:** Akshat Mall, Christopher J. Marx, Jeremy A. Draghi

## Abstract

Heritable non-genetic phenotypic variation—broadly, epigenetics—can potentially influence evolutionary outcomes as direct targets of selection or through interactions with genetic variation. While their evolutionary benefits in generating phenotypic diversity in changing environments is well-characterized, there has been relatively little consideration of how the joint influence of epigenetic changes and mutations would affect traversal of multi-peak adaptive landscapes. Here, we discover general principles for how epigenetics, by generating an epigenetic quasispecies (clusters of semi-stable phenotypes mapped to a single genotype), tends to improve adaptive outcomes of an asexual population on rugged fitness landscapes even without environmental change. In particular, rapid epigenetic changes can sometimes smooth out suboptimal fitness peaks through incorporating fitness contributions of epimutations, allowing access to better adaptive outcomes. Remarkably, the average impact of epigenetics is more strongly influenced by an approximate balance between switching rates rather than the absolute rate at which those switches occur. These findings demonstrate that epigenetic changes can be influential even without having strong heritability and have a striking, yet generally invisible, beneficial role in shaping a population’s adaptive trajectory.

**Significance Statement:** Selection can act upon individuals with epigenetic differences, but it is unclear how much long-term effect this can have on evolutionary trajectories if the epigenetic changes only last a limited number of generations. When the environment changes or more than one functionality is needed simultaneously, it is apparent how bet hedging or division of labour can be advantageous, but what about in a single, constant environment? Here, we find that epigenetics, by allowing individuals rapid yet heritable access to multiple alternate phenotypes, can change the outcome of genetic evolution and has the tendency to remove local fitness peaks and allow adaptation to find higher optima. As such, epigenetics, despite being transient, can profoundly affect adaptive trajectories.

## Introduction

Selection on heritable phenotypic variation is the fundamental driver of adaptation. Epigenetic changes offer a source of heritable phenotypic variation that can partly mimic genetic changes, such as by silencing or activating gene expression, but is distinct in rate and specificity: epigenetic variation occurs orders of magnitude faster than mutation, and targets only specific sites. Genotypes capable of epigenetic switching can produce individuals of not just one, but several distinct, semi-stable phenotypes (“an epigenetic quasispecies”), with their relative frequencies balanced by selection and the rates of epigenetic changes. Typically, the benefits of such epigenetic variation have been modelled only in scenarios necessitating the production of distinct phenotypes, such as the stochastic production of phenotypes suited to possible future environments (bet-hedging)^1,2^, transgenerational plasticity, in response to environmental cues^3^, and division of labour among populations of cells^4,5^. In all of these instances, theory can help us understand how the very high rates of epigenetic switching help to deal with rapid environmental change or otherwise mitigate trade-offs among phenotypes^2,6–10^. However, much less is understood about how the presence of rapidly fluctuating but still heritable epigenetic variation shapes genotype frequencies through quasispecies dynamics during long-term adaptation, even in a static environment.

While existing theory suggests that epigenetics will affect long-term genetic adaptation, the net impact of these effects is hard to predict. Epigenetics, as a source of rapid phenotypic variation, can be a boon for populations at risk of extinction due to rapid environmental change, prolonging survival and potentially interacting with genetic mutations^11–13^. However, epigenetic changes can also compete with mutations, slowing their fixation in the same way that plasticity or behavior can weaken adaptive responses^14,15^. While some theory has focused on the contribution of epigenetic variation in sexual populations with standing genetic variation^16,17^, epigenetic contributions to mutation-limited adaptation are less explored. A few modelling studies have examined epigenetic variation in concert with *de novo* genetic variation, finding that rapid epigenetic variation can sometimes speed evolution^18,19^. The major limitation for interpretation of these studies were their focus upon either a single-peaked fitness landscape, or a scenario where phenotypic effects of genetic and epigenetic changes are completely independent. However, there is abundant evidence that evolution does face rugged adaptive landscapes^20–22^ and it is clear that the same set of loci that are subject to rapid epigenetic switching can also be impacted by beneficial mutations that change gene expression in a similar discrete manner. Here, we simulate the impact of both epigenetic and genetic variation in concert, in populations navigating rugged fitness landscapes to assess when, and how, epigenetics affects evolutionary outcomes.

We hypothesize that epigenetics may help populations better navigate rugged fitness landscapes by avoiding suboptimal peaks. Our primary hypothesis is that rapid epigenetic switching, by generating a cloud of semi-stable “epimutants”, can affect genotype fitness by introducing fitness contributions of several phenotypes, which can smooth out the suboptimal local peaks and valleys that can trap adapting populations. By analogy to viral quasispecies, in which mutations rates are sufficiently high that selection acts on clusters of genotypes in sequence space^23,24^, we propose that epigenetic quasispecies (semi-stable phenotypes mapped to a single genotype) can shape the dynamics of genetic adaptation. While this quasispecies averaging can smooth out constraining features in rugged landscapes, streamlining evolution toward the global peak, rapid epimutation can also have drawbacks. In large populations, evolution can find epistatically favourable combinations of mutations^25,26^; however, rapid epigenetic switching rates could sometimes steer populations down particular trajectories with poor long-term prospects. The instability of epimutations might also frustrate adaptive evolution, eroding the effective fitness of genotypes that are a single epimutation away from a strongly deleterious phenotype. We use simulations and analytical methods to unravel these potential conflicts, identifying when and how epigenetic switching can aid or hinder long-term genetic adaptation.

Here we model asexual, haploid populations evolving on rugged fitness landscapes with both epigenetic and genetic changes. We find that epigenetic switching can help adapting populations better navigate rugged fitness landscapes, specifically aiding them in crossing fitness valleys by smoothing out suboptimal fitness peaks. This benefit is strongly influenced by the fitness contributions from the epigenetic quasispecies for a genotype, which are set by selection and epigenetic transition rates. These results are robust across several orders of magnitude of mutation rates and depend only mildly upon the timescale of epigenetic inheritance in individual lineages of cells.

## Results

### Model development

To understand the effect of epigenetic switching on adaptation in a constant environment, we simulate adaptation with and without epigenetic switching (in addition to genetic changes) on rugged fitness landscapes. We consider a biallelic system of L loci as our genotype space. Here, we limit our analysis to sequences of length L = 5. Each locus contains either the 0 (wild type) or 1 (mutant) allele. We define the genotype with all zeros to be our WT genotype, with a fitness of 1. This WT genotype serves as the starting point of all our simulations. In the absence of epigenetic switching, each genotype maps to a single phenotype, which maps to a single fitness value. The fitness associated with each genotype is defined by the sum of all individual mutation effects and an epistatic term. Genotypes further away from WT, on average, are increasingly fit.

To add epigenetics to simulations of adaptation, we consider the wild-type version of one locus to be capable of epigenetic switching, arbitrarily set as the first locus in this work. This epigenetic locus is capable of stochastic switching between two phenotypic states, which correspond to the WT (0) and mutant (1) states at the locus. Given that microbial examples of epigenetics are typically biallelic and turn genes on and off just like mutations could do, we consider the selective effect of an epigenetic change or a genetic change at that locus to be equivalent. A genetic mutation on the WT allele i.e., 0 → 1, at the locus that allows epigenetic switching breaks the epigenetic switch and locks the mutant phenotype at that locus. Thus, only genotypes with a WT (‘0’) allele in the epigenetic locus can switch epigenetically. As a simplified example, individuals with a genotype “00” can switch between the phenotypic states (and thus fitness) associated with both “00” and “**1**0”. The fitness of an individual thus depends upon on both its genetic and epigenetic state. Epigenetic switching is stochastic with probabilities *r*_*on*_ = 0.01 (from 0→1), and *r*_*off*_=0.1(from 1→0).

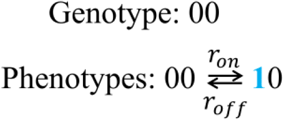

Empirical estimates of rates of epigenetic switching are sparse and vary between systems, but are considerably higher than mutation rates^7,8^. Epigenetic switching rates are often observed to be asymmetric, with switching towards one phenotype being faster than the alternate. In the *pap* operon in *E. coli*, the switching rates from an On to Off state is ∼100-fold higher than the Off to On state^27^, with the individual rates estimated to be around 10^−2^ and 10^−4^respectively, per generation. In the case of *E. coli* persisters, switching rates are estimated to be 10^−6^into a persister phenotype, and 10^−1^out of a persister phenotype^28^.

Mutations arise with a genome-wide rate μ = 10^−6^and are bidirectional; in most organisms, mutation rates at a particular target tend to be ∼ 10^−6^−10^−8^per generation^8^. The effect of a mutation on the individual depends on its phenotype, which depends on both the genetic and epigenetic state of the individual. We use a discrete-time Wright Fisher model to simulate adaptation with a fixed population size N = 10^5^, allowing both genetic and epigenetic changes. Organisms are haploid and reproduce asexually.

### Epigenetics can help reach more fit peaks on rugged fitness landscapes

We first studied the impact of epigenetics on adaptation as a function of ruggedness of the underlying fitness landscape. We generated 1000 different fitness landscapes, encompassing a range of ruggedness. For most parameter combinations surveyed, almost all populations eventually end up on one of the fitness peaks within 10^4^generations. For each landscape, we quantify which of the available peaks in the landscape the population ends up on (across 200 replicate simulations). To quantify the optimality of adaptive outcomes, we define a parameter “rank” which is the relative position of the peak reached by the population among all possible peaks; the most fit peak in the landscape is assigned rank 1, the second most fit peak is assigned rank 2, and so on. We classify these landscapes by ruggedness, defined as the number of fitness peaks, and record the average rank of final fitness peak reached by adapting populations with and without epigenetics as a function of ruggedness. Figure 1A shows that epigenetic switching (G+Epi), on average, aids the population in reaching more fit peaks than with only genetic mutations (G only), with the benefit increasing with the ruggedness of the landscape. As the number of peaks grows past one, populations are more likely to get stuck at suboptimal fitness peaks instead of reaching the global peak, and the average rank of final peak reached increases. However, epigenetic switching, on average, assists adapting populations in reaching more fit peaks, a pattern which is similar if we use alternate measures of adaptive outcomes like average fitness, or probability of reaching the global peak (Fig S1); or alternate ways of constructing fitness landscapes like the house of cards model (Fig S2).

**Fig 1.**
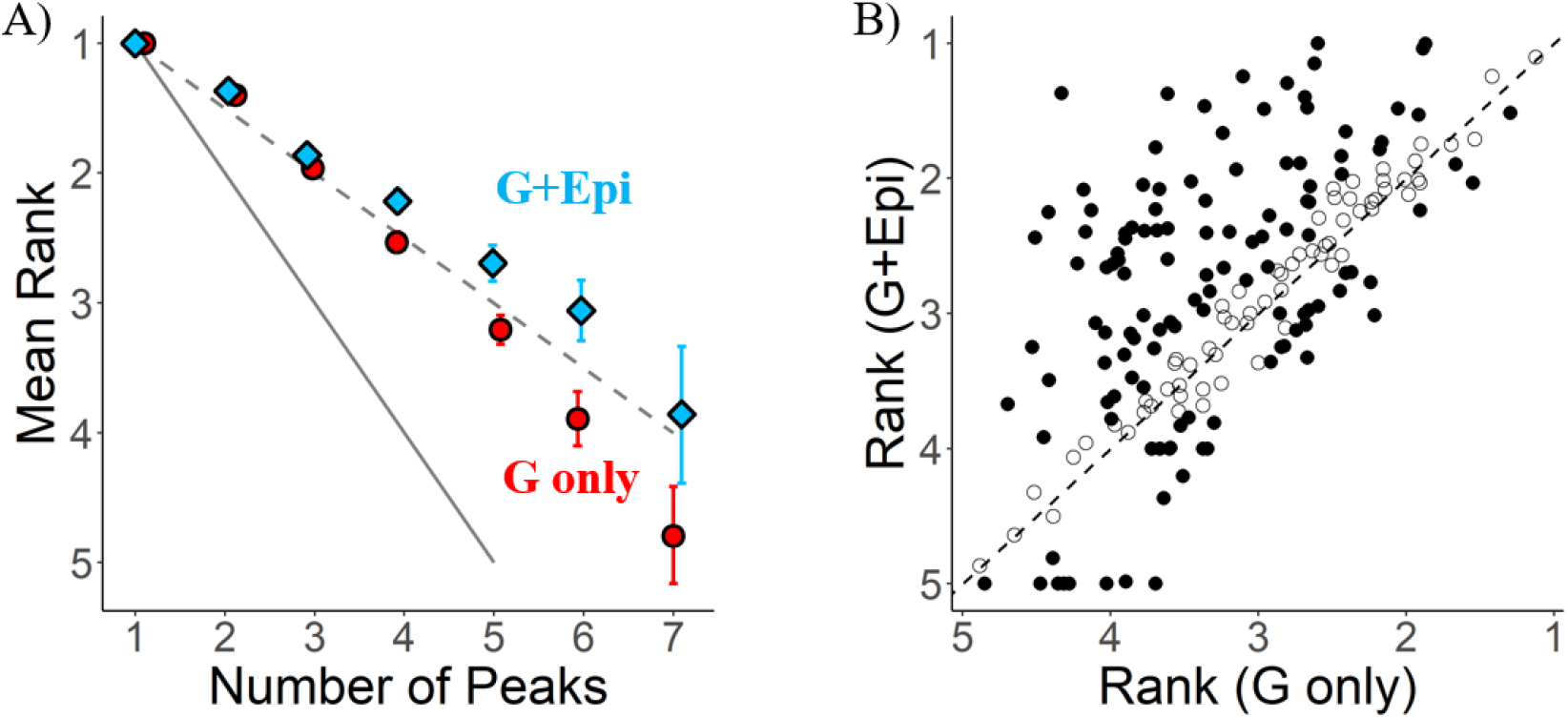
A) Epigenetics helps reach more fit peaks in rugged landscapes. Each point denotes average rank of final peak reached for all landscapes of a particular ruggedness (G only=genetics only, G+Epi=genetics + epigenetics). Error bars denote 95% confidence intervals calculated by bootstrapping. Dashed line indicates expected trend when all available peaks are reached with equal probability. Solid line indicates maximum possible rank for a given number of peaks. **B) Heterogeneity in effect of epigenetics on final rank for each 5-peaked landscape**. Each point represents the average final rank (after 200 replicate simulations) on one landscape, with (G+Epi) and without (G only) epigenetics. Dashed line represents x=y. Points above the line represent landscapes where epigenetics had a positive effect – final rank was lower with epigenetics than without. Points below the line represents landscapes where epigenetics had a detrimental effect. Points are filled when the mean rank with and without epigenetics are significantly different (p<0.01, Welch’s t-test).

To ask how consistently epigenetic variation helps populations reach higher peaks, we focus on outcomes among a subset of 189 landscapes that all have five peaks, and therefore similar degrees of ruggedness. In Fig. 1B, we observe that epigenetic switching can have a range of effects on the final rank – beneficial (solid circles above x=y line, 48%), statistically neutral (unfilled circles near x=y line, 37%), and even detrimental (solid circles below x=y line, 15%). This emphasizes a critical feature – while we can infer statistical patterns about the average impact of epigenetics on adaptation (Fig 1A), the effect on any particular landscape can be strongly contingent upon the idiosyncratic structure of that fitness landscape (Fig 1B).

### Epigenetic switching can help cross fitness valleys

We hypothesized that epigenetic quasispecies, by incorporating fitness contributions from alternate phenotypes, can reduce or eliminate fitness valleys trapping populations at suboptimal peaks, and aid adaptation on rugged fitness landscapes. Adopting the basic dynamics of mutation-selection balance, we derive a simple expression (Box 1) to illustrate the hypothesized role of epigenetic quasispecies helping reduce the constraint of fitness valleys. When epigenetic changes are possible (i.e., that genotype has not acquired a mutation that breaks epigenetic switching at that locus), each individual genotype does not map to a single phenotype, but to an epigenetic quasispecies of two alternate phenotypes, each with their associated fitness. Given the high rates of epigenetic switching, the “effective fitness” of a genotype equilibrates to a weighted average that depends upon the relative frequencies and fitness of the alternate epigenetic states. For a genotype, the relative frequency of each epigenetic state is dictated by the switching rates, and selection. We calculate the equilibrium frequency of each epigenetic state as a function of switching rates, and fitness’ of alternate epigenetic states (Eq. 2). We can then use the equilibrium frequency to calculate the effective fitness of a genotype as the weighted sum of individual fitness contributions from each epigenetic state (Eq. 3).

#### Box 1

Consider a genotype expressing two alternate epigenetic states *P* and *Q*, switching at rates *a* and *b*.

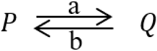

Fitnesses of *P* and *Q* are 1 and 1 + *s*,respectively. At generation *i*, frequencies = *p*_*i*_ and *q*_*i*_. (*p*_*i*_ + *q*_*i*_ = 1)The relative frequencies of *P* and *Q* depend on the switching rates and their relative fitness. In generation *i* + 1, the new frequency of epigenetic state *P* can be estimated as:

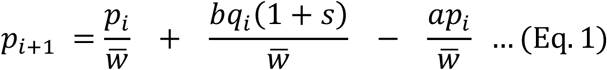

Where mean fitness is 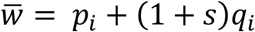 . Selection and switching will drive the alternate phenotypes towards an equilibrium where *p*_*i*+1_ = *p*_*i*_ = *p*^∗^ (say). We derive an expression whose solution gives us the equilibrium frequency of epigenetic state *P* :

*sp*^∗2^ + *p*^∗^[(1 −*a*)−(1 + *s*)(1 + *b* )] + *b* (1 + *s*)= 0 … (Eq. 2)Now, *q*^∗^ = 1 −*p*^∗^. The fitness of the genotype can now be re-defined as *p*^∗^(1)+ *q*^∗^(1 + *s*)… (Eq. 3)

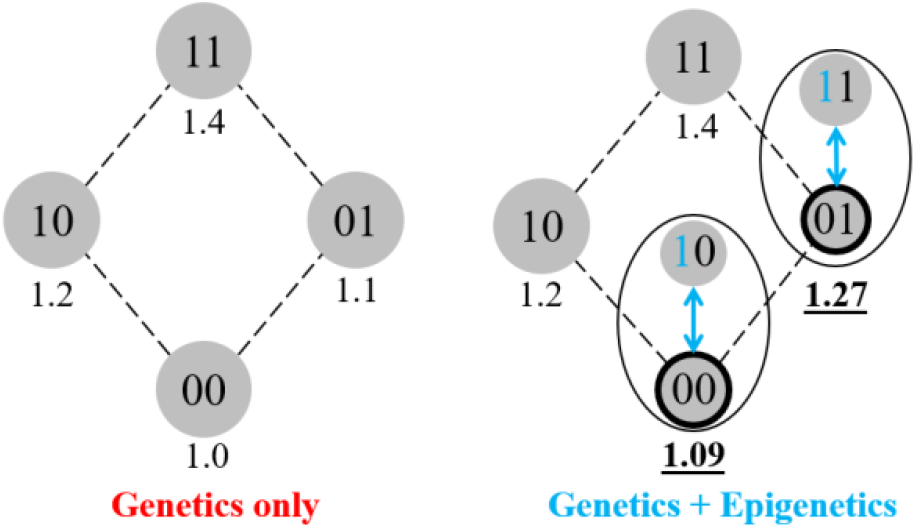

In this simplified illustration, grey circles represent genotypes, with the sequence represented inside, and fitness below the circles. Genotypes one mutational step away from each other are connected by black lines. When epigenetic switching is allowed at the first locus, genotypes with a WT (“0”) allele in the first locus (nodes outlined in black), allow access to alternate phenotypes associated with the mutant (“1”) allele in the first locus (connected by double-sided blue arrows). The genotype fitness is now a function of the epigenetic quasispecies as a whole, and can be re-evaluated as shown above.

To demonstrate that accounting for the fitness contributions of epimutants leads to a decrease in the effective number of local peaks in a rugged landscape, we used the approach in Box 1, and calculated effective fitness for each genotype capable of epigenetic variation among the same set of 5-peak landscapes. We re-evaluated the number of peaks in the landscape, and based on these calculated fitness values, find that epigenetics reduces the effective number of peaks in 70% of the landscapes, often removing several suboptimal peaks (Fig S3). We quantified a measure “rank improvement”, which is the difference in average final rank for a landscape with and without epigenetic switching allowed. A positive value indicates that adaptation with epigenetics reached more fit peaks, while a negative value indicates adaptation with epigenetics reached less fit peaks compared to purely genetic adaptation. This improvement correlates with the number of peaks removed by calculating effective fitness (from Box 1) with epigenetics (Figure 2). For landscapes where epigenetic switching eradicates some local fitness peaks, average final rank improves for adaptation with epigenetics, in strong concert with the number of local fitness peaks removed. The number of peaks removed explains a substantial proportion of the variance in adaptive outcomes with and without epigenetic variation (*R*^2^ = 0.31).

**Fig 2.**
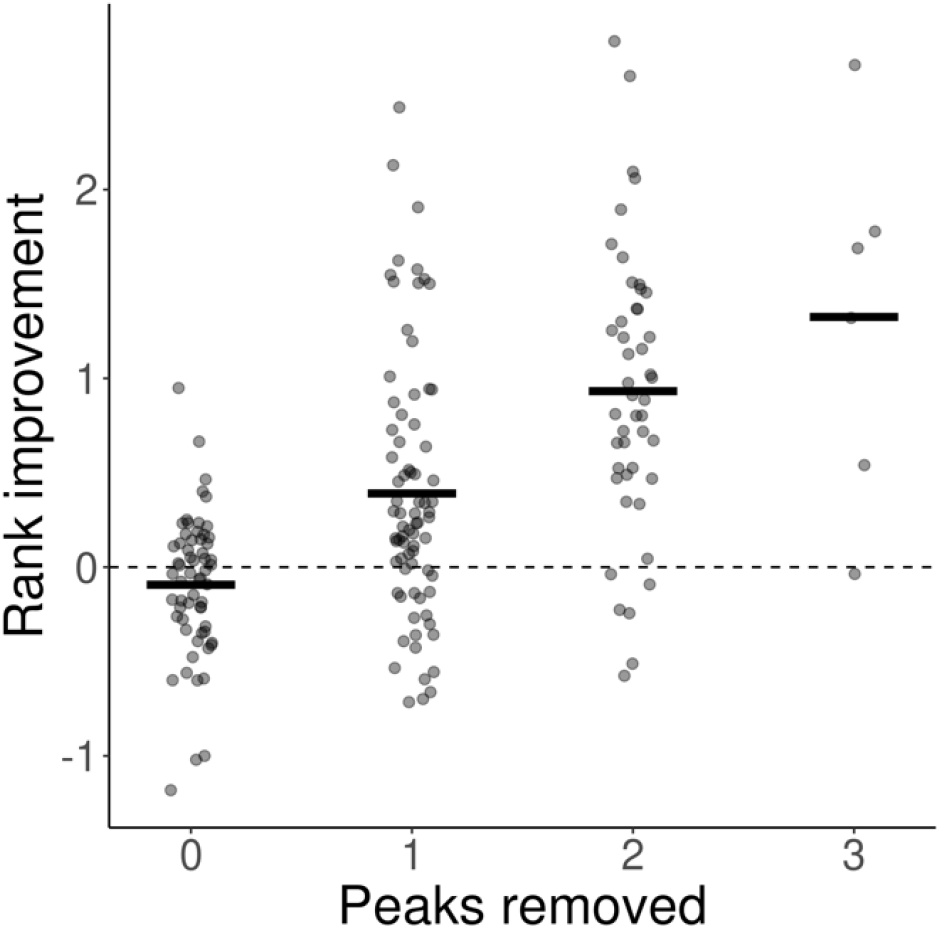
Epigenetics tends to result in better outcomes when it erases local peaks. For each 5-peak landscape, the improvement in average final rank with epigenetics is plotted on the y-axis. Using the equations in Box 1, we can replace the fitness of genotypes capable of an epimutation with a new effective fitness. Landscapes are classified by how many local peaks are removed in these new calculated landscapes.

To illustrate how epigenetic switching can help populations cross fitness valleys, we focus on one example landscape with five fitness peaks. All five peaks can be accessed during genetic evolution, though many populations remain trapped at the worst one and very few reach the global peak (Fig 3A). Enabling epigenetic switching has complex effects on the frequencies of each peak among our ensemble of replicate populations, but generally improves the outcome and prevents populations from being stuck at the lowest peak (Fig 3B).

**Fig 3.**
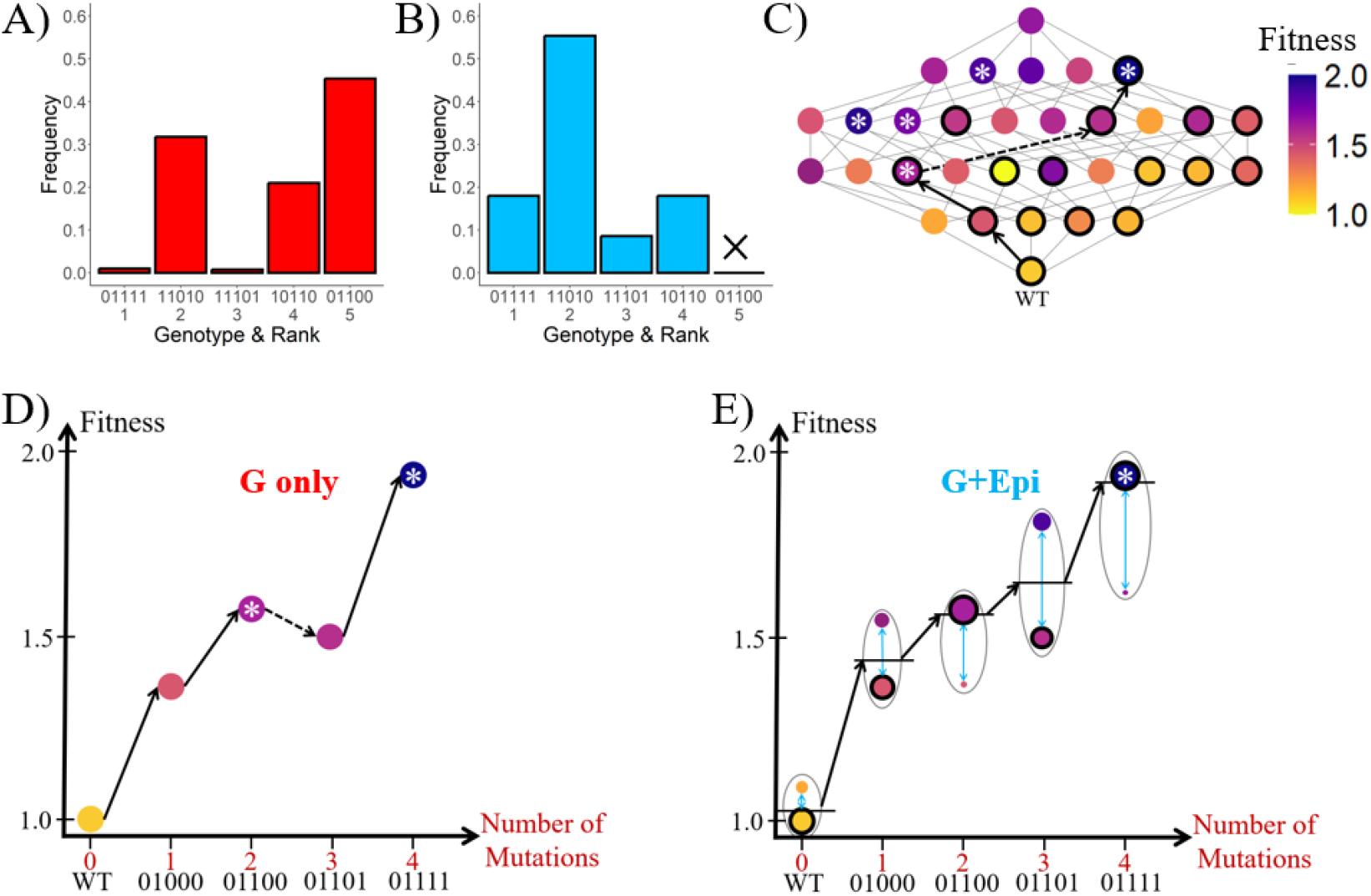
Epigenetics can improve adaptive outcomes by removing suboptimal peaks. **A)** Probability of reaching each peak with genetic changes only. Here, suboptimal local peaks trap most populations. **B)** Probability of reaching each peak with genetic + epigenetic changes. The probabilities are significantly different (*p* < 10^−15^, Pearson’s chi-squared test) from the ones associated with genetic changes only, with increased probability of populations ending up on peaks of higher fitness. The 01100 genotype is eradicated as a peak. **C)** Example fitness landscape and an adaptive trajectory we focus on in the landscape. Each node represents a genotype. Colors denote fitness, with darker shades for higher fitness. Genotypes one mutation away from each other are connected by gray lines. Genotypes capable of epigenetic switching denoted by a black outline. Fitness peaks are denoted by asterisks *. This particular landscape houses 5 local fitness peaks. We focus on the adaptive path connected by dark black lines in D and E. **D)** The adaptive steps associated with the trajectory with only genetic changes allowed. The population gets stuck at a suboptimal peak (“01100”) and further adaptation requires a deleterious step (dashed arrow). **E)** The adaptive steps associated with the trajectory with genetic & epigenetic changes allowed. With epigenetic changes allowed at the first locus, individuals can access alternate phenotypes (new circles connected by bidirectional blue arrow). Alternate epigenetic states (epigenetic quasispecies) of the genotypes are represented by circles connected with the genotypes by double-sided blue arrows, and the pair of epigenetic states encased by a bubble. Relative size of circles in the bubble represent the relative equilibrium frequencies of the alternate epigenetic states. The effective fitness of the genotype (shown by solid horizontal black line across the available phenotypes) is a weighted sum of alternate phenotypes (Box 1). In such a regime, the previously deleterious step (dashed arrow in 3C) is now beneficial (solid arrow here).

To understand the root cause of these beneficial effects, we focus on one landscape from our set of 5-peaked landscapes used, and illustrate an adaptive path (Fig 3C) leading to both the worst local peak and the global optimum. With only genetic adaptation, the fitness steps associated with each step in the trajectory are represented in Fig 3D. The population acquires two successive beneficial mutations and reaches a local fitness peak “01100”, from which all one-step mutations are deleterious (dashed arrow); the population is now temporarily stuck at the suboptimal peak. When epigenetic switching is allowed at the first locus, adaptive outcomes along the same initial trajectory change (Fig 3E). The epigenetic quasispecies behavior is illustrated by the difference between Fig. 3 D & E—replacing each fitness with a weighted equilibrium fitness value removes the suboptimal peak, allowing evolution to smoothly progress towards the global peak. Without epigenetics, evolution would need to cross this fitness valley with a long waiting time at realistic mutation rates^29^. At the end of adaptation, the population harbours no signature of epigenetic switching (since any epigenetic states revert back to the WT phenotype and the epigenetic locus is retained as the WT allele throughout). Yet, epigenetic switching was critical to efficiently navigating the fitness landscape and shaping adaptive outcome.

### Epigenetics can aid valley crossing across a range of mutation and switching rates

Adaptive outcomes on epistatic fitness landscapes depend on mutation rates, and potentially epigenetic switching rates. To explore how changing mutation or switching rates impact our observation of epigenetics helping cross fitness valleys and leading to more fit peaks on average, we simulated adaptation with and without epigenetics on the same set of 5-peaked landscapes as earlier, with changing values for either mutation or switching rates. We first focus on changing mutation rate.

We observed that epigenetics helped reach more fit peaks across a range of mutation rates (Fig. 4A). At low mutation rates (μ∼10^−7^ −10^−5^, or Nμ < 1), there is little effect of increasing mutation rate for the case of purely genetic adaptation. However, there is a modest benefit of increasing mutation rate on adaptation with epigenetic switching because in this regime, adaptation is mutation-limited. While epigenetics helps reduce or eliminate fitness valleys, mutations are still needed to cross these diminished valleys, and increasing mutation rate leads to an increase in the instances of valley crossing, which allows populations to reach more fit peaks within the simulated period. For high mutation rates (μ > 10^−5^), adaptive outcomes improve sharply for both cases, with and without epigenetic switching; and slowly converge with further increase in mutation rate. This is because genetic changes now occur rapidly enough and populations can escape local peaks via genetic changes alone^29^. This leads to a sharp decrease in average final rank for both scenarios, and the relative benefit conferred by epigenetics diminishes. We observe the same pattern when we use alternate measures of quantifying adaptive outcomes like mean fitness, or probability of reaching global peak (Fig S4).

**Fig 4.**
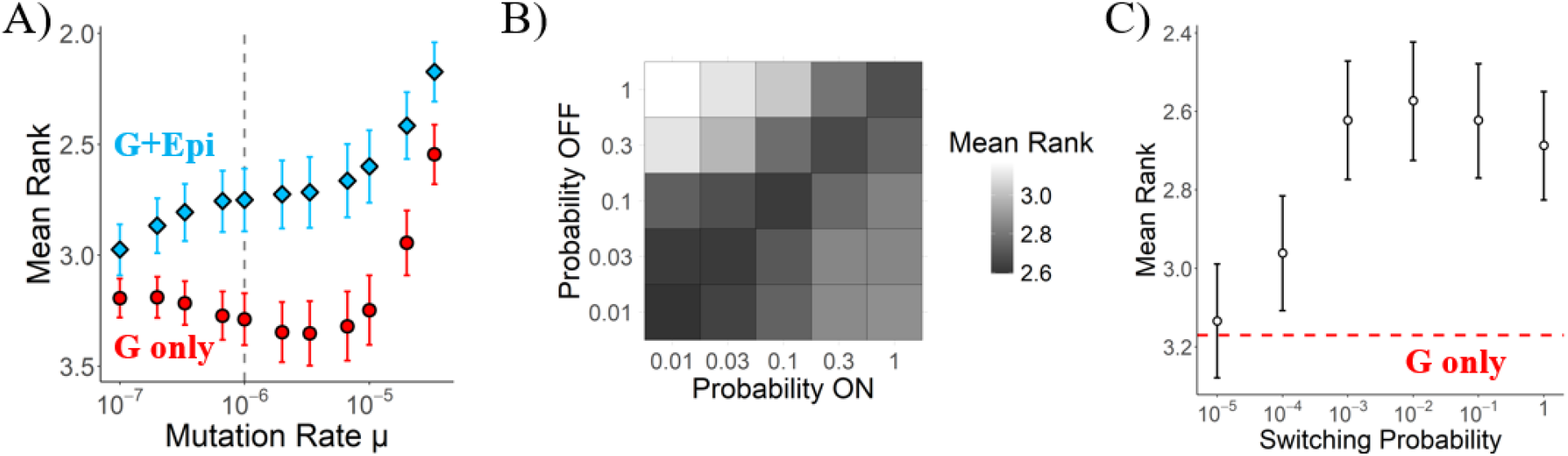
Epigenetics is beneficial over a wide range of mutation and epigenetic switching rates. **A)** Mean Rank vs Mutation rate with (G+Epi) and without (G only) epigenetics. Each point represents the average final rank over all 5-peaked landscapes. Error bars denote 95% confidence intervals estimated by bootstrapping. Dashed line represents mutation rate considered for all results above. **B)** Mean rank for combinations of epigenetic switching probabilities between 0.01 and 1. Darker colors represent lower average rank i.e., better adaptive outcomes. **C)** Mean rank vs epigenetic switching probability (*r* = *r*_*on*_ = *r*_*off*_). Each point represents the average final rank over all 5-peaked landscapes. Error bars denote 95% confidence intervals estimated by bootstrapping. Red dashed line represents the average final rank for purely genetic adaptation.

We next focused on the impact of epigenetic switching rates, specifically how the ratio of rates between the two directions affects adaptive outcomes. We find that a balance between rates led to better adaptive outcomes (Fig 4B, and Fig S5), relative to the rates being strongly biased in either direction. To understand how the absolute magnitude of epigenetic switching rates shapes adaptive outcomes, we estimated average final rank of populations adapting with and without epigenetics across multiple orders of switching rates (Fig 4C). The absolute rates generate a tension between two opposite effects of epigenetics upon adaptation: high rates of epigenetic switching make it more advantageous compared to mutation, yet the same high rates also create lineages that stay in that phenotypic state for less time. For simplicity, we focused on symmetric epigenetic switching rates i.e., equal rates in both directions. We observed that epigenetics helps adapting populations reach more fit peaks over a wide range of switching rates (points above red dashed line in Fig. 4C). At very low switching rates (≤ 10^−5^), adaptive outcomes are largely the same as with purely genetic adaptation (red dashed line in Fig 4C), as the impact of epigenetics at those rates are no different from that of genetic changes. With increasing switching rates, the benefit of epigenetics increases, reaching an optimum. A further increase in switching rates leads to a slight increase in average final rank as individuals with extremely high switching rates incur a cost of diminished visibility to selection, even if more fit. Nevertheless, even high switching rates fare significantly better than purely genetic adaptation, indicating a wide window for epigenetics to be beneficial and aid populations reach more fit peaks. We observe the same pattern for alternate measures of quantifying adaptive outcomes (Fig S6) and in analysis of individual landscapes that had distinct responses to epigenetics (Fig S7).

## Discussion

In this work, we show that epigenetics, via generation of epigenetic quasispecies which diminish the constraint of fitness valleys, allow adapting populations to better navigate rugged fitness landscapes to reach more fit peaks (Fig. 1A). The influence of epigenetics is complex, hindering evolution on some landscapes, having little effect in others, and greatly boosting the rank of the final peak in many (Fig. 1B). Despite this heterogeneity, we found that we can understand much of the effect of epigenetics on evolvability by calculating an effective genotypic fitness as an equilibrium of the epigenetic quasispecies as a whole (two alternate phenotypes in this work, Box 1), and use this to calculate how many suboptimal peaks were eradicated (Figs. 2 & 3). These equilibrium calculations also explain why epigenetics can aid evolvability even at very rapid switching rates, which prevent epimutations from fixing as actual steps in an adaptive trajectory (Fig. 3B).

The results here demonstrate a novel mechanism for valley crossing in asexual populations. The idea that rugged fitness landscapes constrain and complicate evolution is nearly a hundred years old (reviewed in^30^), and theoretical studies have thus far identified a number of ways adapting populations can cross fitness valleys^29,31–36^. Mechanisms of speeding valley-crossing can be separated into those that help populations move better through fitness landscapes (e.g., high mutation rate) or those that work to compress peaks and valleys, smoothing out rugged landscapes. Examples of the latter class include high-frequency but not heritable variation due to stochastic variability in phenotypes^37–39^. In this light, epigenetic variation that occurs rapidly and is heritable combines both ideas, aiding movement across fitness valleys and also diminishing them when fitness is an average of a cloud of closely related phenotypes. Thus, our work bridges models of very high genetic mutation rate, such as viral quasispecies, with models of purely non-heritable phenotypic heterogeneity^37^. As seen in Figure 4B, the biggest benefit of epigenetic variation is found at intermediate levels of heritability, supporting a uniquely useful role for this kind of variability.

While any given epigenetic locus may have arisen due to selection to generate distinct phenotypes for alternate environments or roles^40–44^, the consequence of epigenetics is felt even during adaptation to static environments. While our model differs from many previous approaches to epigenetics by focusing on an unchanging environment, it will be interesting to explore the interaction between the effect of epigenetic quasispecies on genetic adaptation in concert with environmental change. One clear place for exploration is to allow epigenetic switching rates to be dependent upon the environment. This can aid rapid adaptation without incurring a strong fitness cost in alternate environments^45^, and allow for persistence of “epigenetic memory” of past environments^46^, as rates increase transiently only in one direction, as is known to be the case in many systems, such as the *pap* operon involved in production of pili in uropathogenic *E. coli*^47,48^. The fact that clinical isolates have been found to exhibit increased switching rates, while still environment-dependent, compared to lab strains^49^ suggests recent selection on epigenetic switching rates. Future work could combine our fitness-landscape approach taken here with epimutations whose rate depends on the organism’s environment to understand both initial and longer-term adaptive consequences.

Epigenetic variation is widespread across all domains of life, suggesting that the effects described here could be quite general. Empirical examples of epigenetic switching are known via mechanisms such as heritable changes in DNA methylation^27,50^, multistable metabolic networks^51,52^, and others^45,53^, often regulating critical phenotypes such as virulence in *Salmonella*^54^. Instances of long-term adaptation shaped by epigenetic changes have also been observed, in evolution of cancers^55^, speciation events^56^, and pathogen adaptation^57^. Although characterized examples of microbial epigenetics are biased towards pathogens and virulence traits^58^, instances where methylomes have been characterized for other bacteria, dozens of genes have been discovered to be under epigenetic regulation via alternate DNA methylation^58,59^. Since pathogens like *Salmonella* have already been characterized to have cell-to-cell heterogeneity in expression of 10 loci under epigenetic regulation^60^, this suggests a massive number of semi-stable phenotypes may be accessible from a single genotype (e.g., for just ten loci, 2^10^ = 1024alternate phenotypes). This number is likely higher when one considers the prospect of epigenetic change via alternate mechanisms. Our model represents a minimal version of an epigenetic quasispecies with only one locus capable of epigenetic changes, but further work can explore how the impact of epigenetics scales with increasing number of loci capable of epigenetic switching.

As the theorized roles of epigenetics in shaping adaptation become stronger, empirical work needs to stringently consider the possibility of epigenetic changes influencing long-term genetic adaptation. Most studies of evolution in lab or in nature have traditionally focused on genetic changes alone, and adaptive phenotypes not explained by beneficial mutations remain underexplored. This is partly because of an underappreciated role of epigenetic changes in shaping long-term adaptation, but also because of the difficulty in detecting such changes.

However, recent advances allow some non-genetic factors, such as the methylome, to studied in great resolution. Studies employing these techniques have discovered key and widespread roles for epigenetic changes, like in gut microbiome evolution^61^, pathogen evolution^57^, and cancer evolution^62,63^. Relatedly, the advent of single-cell RNA-seq for microbes has shown that nearly all environments have distinct clusters of cell phenotypes originating from a single genotype^64^ which, whether or not they are caused by alternate epigenetic states, could have their fitness represented in a similar manner as a quasispecies of expression types. These empirical findings, and our results in this work, indicate a strong potential role of epigenetics, despite its transience, in shaping adaptive outcomes. Increased attempts to characterize non-genetic factors during the course of evolution would shed further light on the role of epigenetic changes in driving and shaping the course of evolution.

## Materials & Methods

### Constructing fitness landscapes

We define a genotype σ^∗^ with all zeroes to be our WT genotype, with a fitness of 1. This genotype serves as the starting point of all our adaptive walks. All other 2^*L*^ −1 genotypes have one or more mutations relative to the WT. All single-step mutants from WT are assigned a fitness 1 + *s*_*i*_, with *s*_*i*_ drawn from an exponential distribution with mean 0.2, and fitness for other genotypes σ is defined the following manner – *f*(σ)= *Σs*_*i*_ + η(σ). *Σs*_*i*_ represents the sum of individual mutation effects. η(σ) represents an epistatic term, and is a random number drawn from a normal distribution with mean = 0, and standard deviation = 0.25. Using this approach, we generate 1000 fitness landscapes, which stochastically differ in their ruggedness with number of peaks ranging from 1 (completely smooth) to 7.

### Simulating adaptation on fitness landscapes

To simulate adaptation, we use a Wright Fisher model, with a fixed population size = N. At each timestep, N individuals are selected for the next generation based on probabilities weighted by fitness. These chosen individuals are then allowed to mutate and change epigenetic states with their respective probabilities. This process is iterated for 10,000 generations. The key parameters for simulations are listed below & are held constant for all simulations, unless otherwise mentioned in the main text.

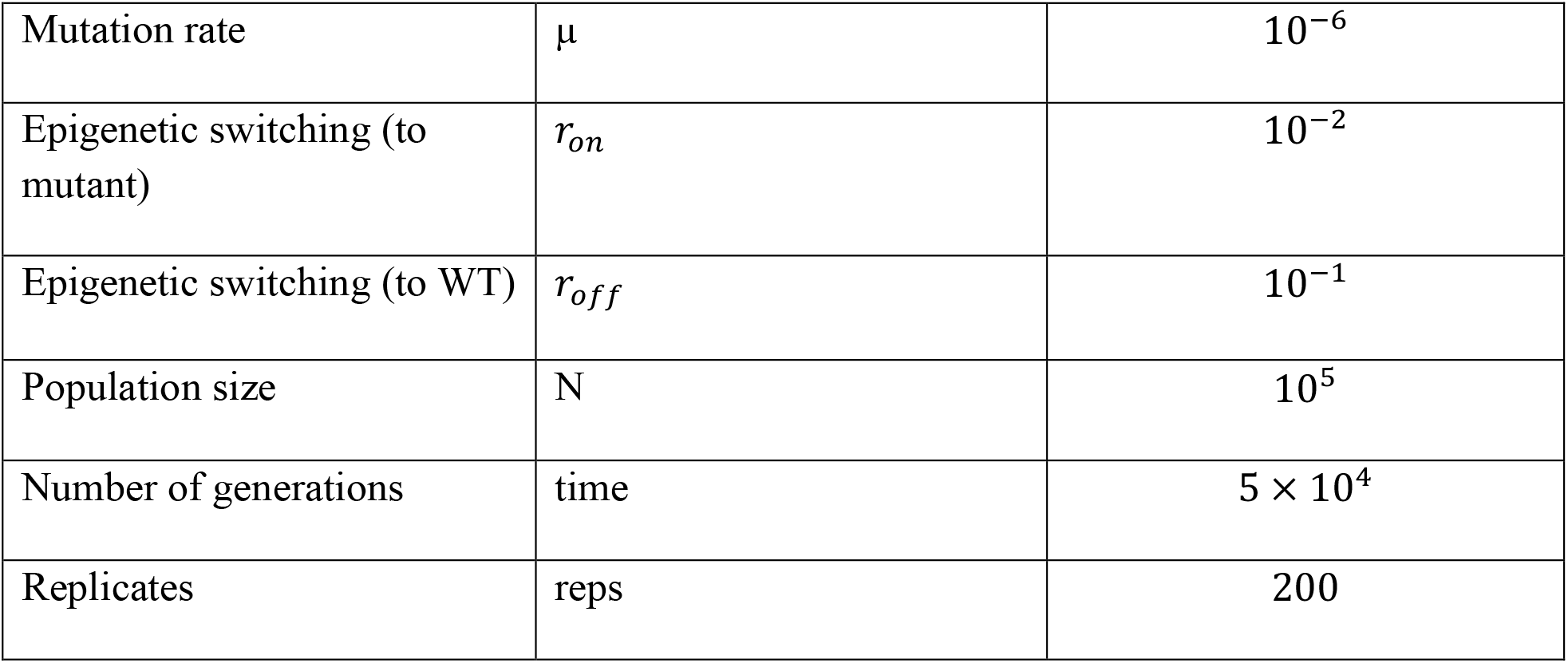

## Data Availability

The code and data for this work are available on Github. https://github.com/mallakshat/EpigeneticQuasispecies

## Acknowledgements

We are grateful to Daniel Segrè, C. Brandon Ogbunu, Daniel Schultz, Matthew Kinahan, Mohammad Abbaspour, and Noah Arts for helpful feedback on the manuscript; to Alexander Alleman for discussions; and members of Marx-Udekwu and Draghi labs. This research was supported by US Department of Energy grant DE-SC0022318 to CJM and JAD.

## Supplementary Information

**Fig S1.**
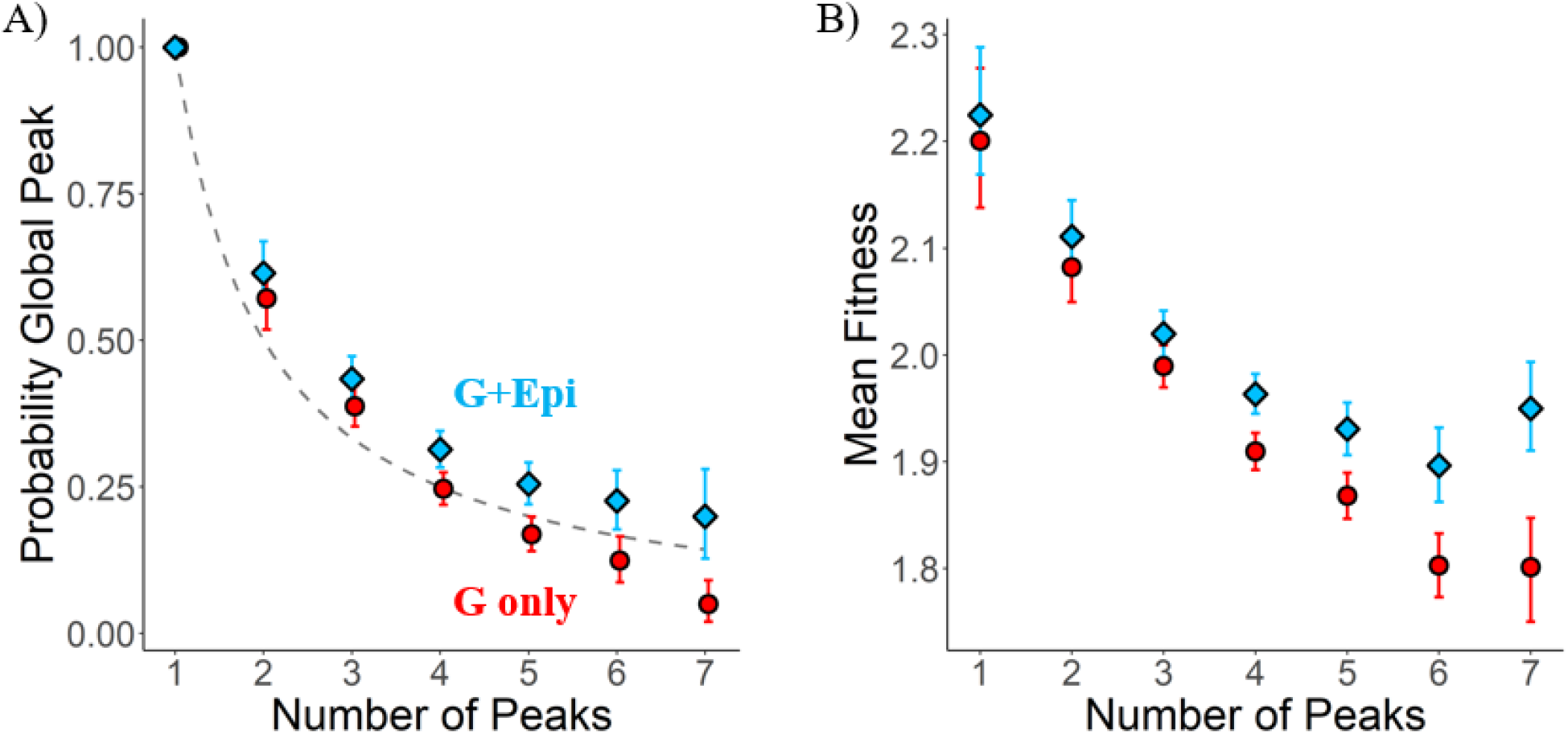
Epigenetics helps better navigate rugged fitness landscapes. We use alternate measures – **A) probability of reaching global peak**, and **B) average fitness** to contrast to the average rank measure used in Fig 1A. in the main text. In both panels here, each point depicts the average of the measure on the y-axis for all landscapes of a particular ruggedness (G only = genetics only, G+Epi = genetics + epigenetics). Error bars denote 95% confidence intervals calculated by bootstrapping. In A) Dashed line indicates expected trend where probability of reaching global peak is simply = 1/number of peaks.

**Fig S2.**
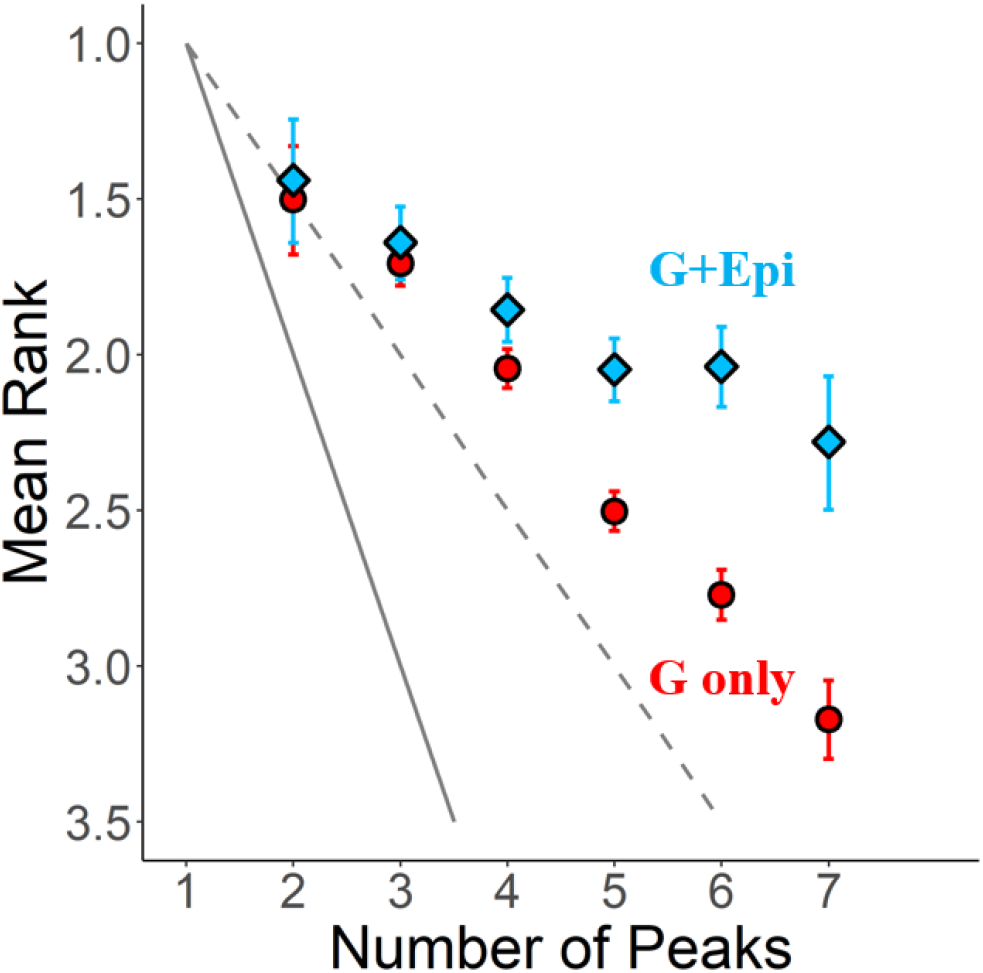
Epigenetics helps reach more fit peaks for alternate models of constructing landscapes. Here, we use a House of Cards model to construct uncorrelated fitness landscapes. Each point represents the average rank of final peak reached for all landscapes of a particular ruggedness (G only = genetic alone, G+Epi = genetics + epigenetics). Dashed line represents expected trend if all local peaks were reached with equal probability. Solid line represents the minimum rank possible for a given number of peaks. Error bars denote 95% confidence intervals estimated by bootstrapping.

To construct fitness landscapes following the House of Cards (HoC) model, we define a genotype σ^∗^ with all zeroes to be our WT genotype, with a fitness of 1. This genotype serves as the starting point of all our adaptive walks. All (2^*L*^ −1) mutant genotypes are assigned a fitness randomly drawn from a normal distribution with mean = 1, and standard deviation = 0.25. Using this approach, we generate 1000 fitness landscapes, which stochastically differ in their ruggedness with number of peaks ranging from 2 to 10. We do not obtain any single-peaked landscape using this approach.

**Fig S3.**
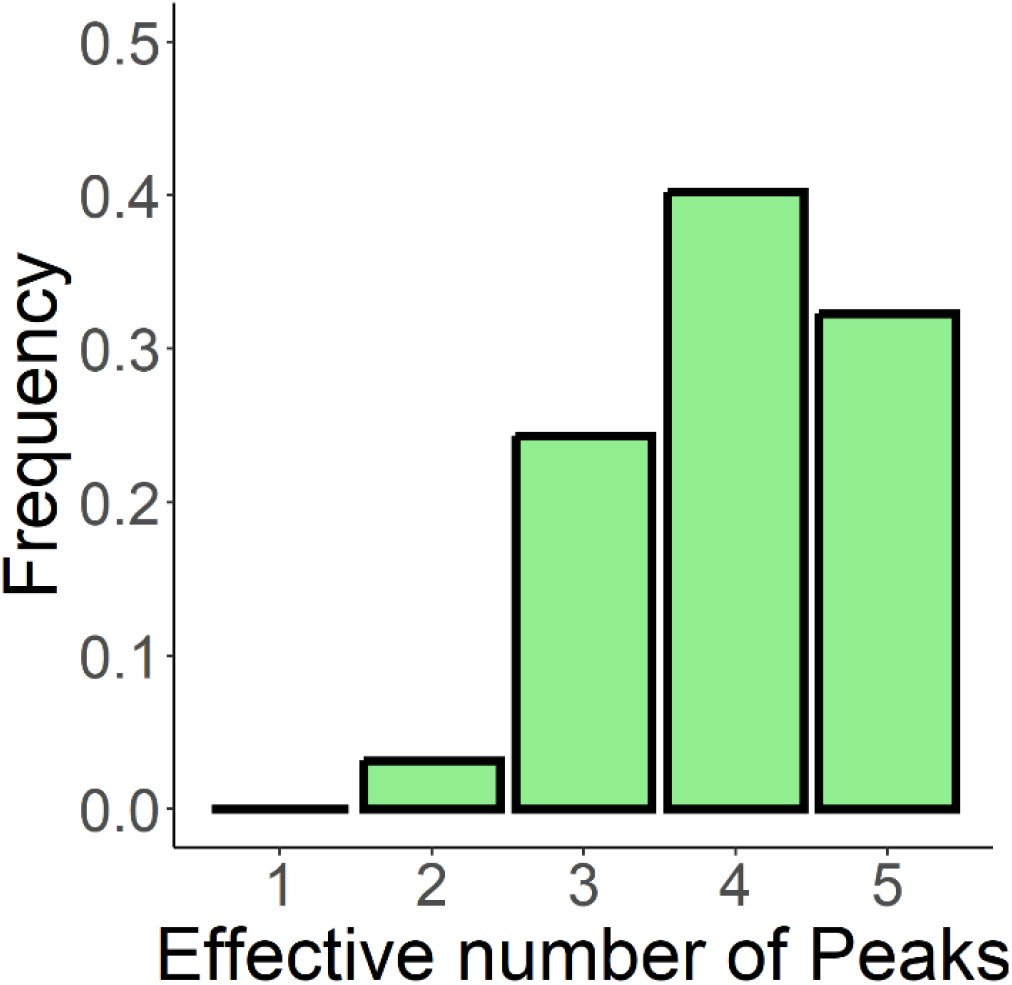
Epigenetics helps reduce the number of local peaks.

We analyse (using Box 1) the effective fitness of a genotype incorporating fitness contribution of the epigenetic quasispecies as a whole, and estimate the new number of local peaks for the set of 5-peaked landscapes used in the main text. For ∼30% of the landscapes, we see no change in the effective number of peaks, whereas epigenetics helps eradicate suboptimal local peaks for ∼70% of the landscapes.

**Fig S4.**
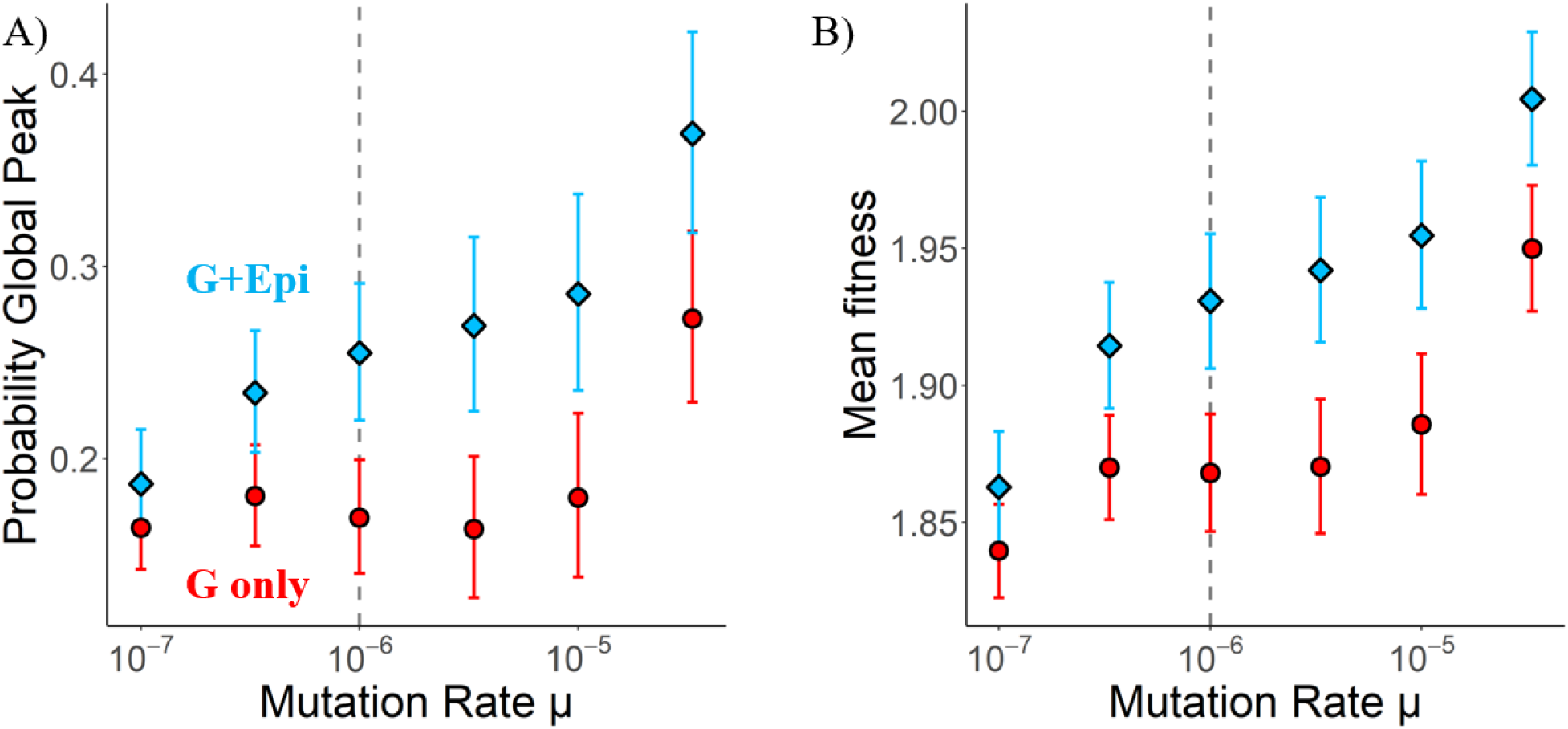
Epigenetics is beneficial over a wide range of mutation rates. We use alternate measures – **A) probability of reaching global peak**, and **B) average fitness** to contrast to the average rank measure used in Fig 4A. in the main text. In both panels here, each point depicts the average of the measure on the y-axis for all 5-peaked landscapes for a given mutation rate (G only = genetics only, G+Epi = genetics + epigenetics). Error bars denote 95% confidence intervals calculated by bootstrapping. Dashed vertical line indicates the mutation rate considered for all other results in the main text.

**Fig S5.**
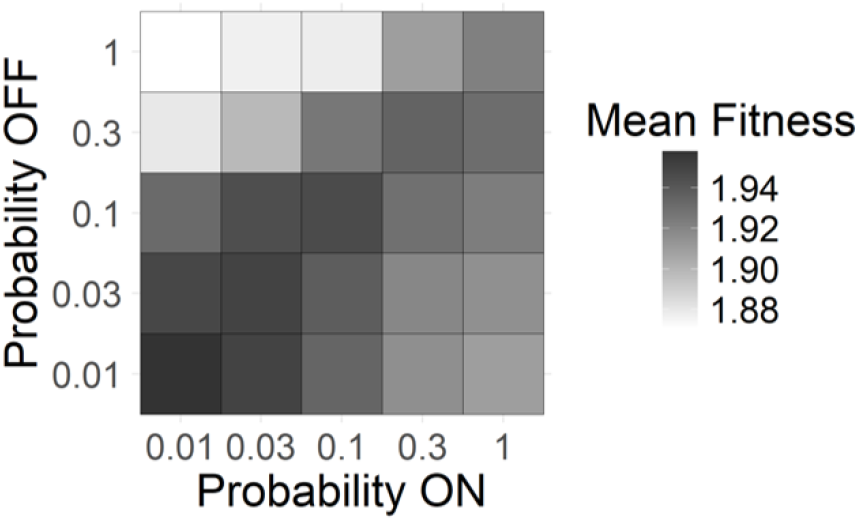
Epigenetics is beneficial for balanced switching probabilities. Mean final fitness (over all 5-peaked landscapes) for epigenetic switching probabilities between 0.01 and 1. Darker colors represent more fit outcomes. Same experiment as Fig 4B in main text but fitness used as a measure instead of final rank.

**Fig S6.**
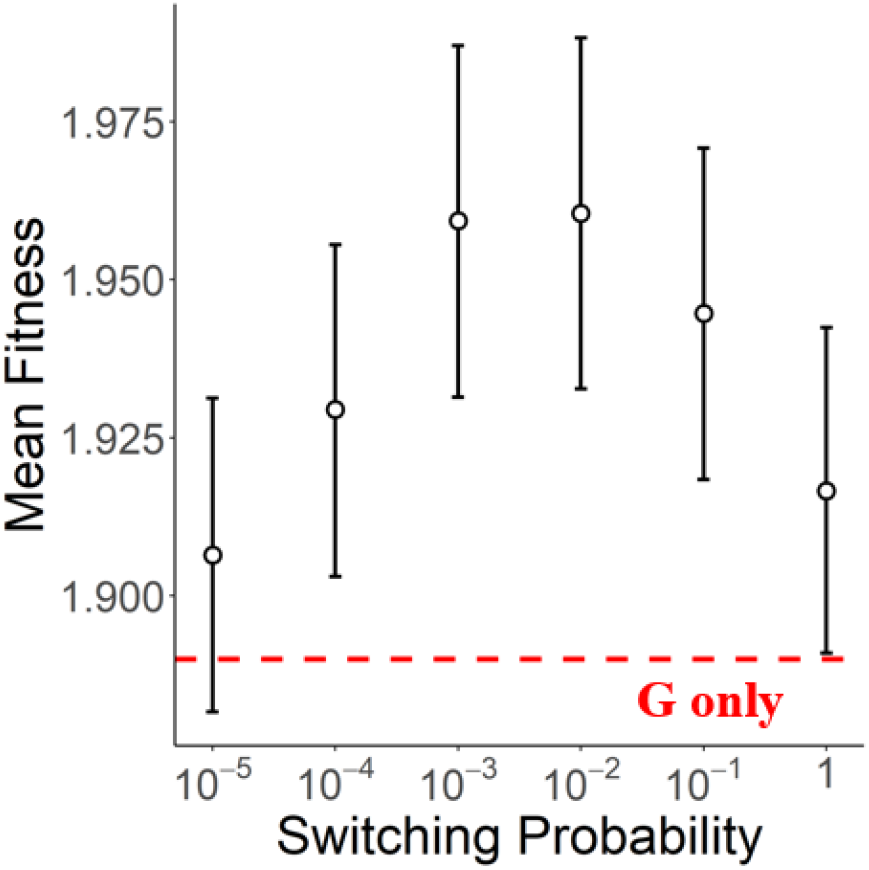
Intermediate epigenetic switching probabilities are optimal. Each point represents average final fitness over all 5-peaked landscapes, for a specific value of epigenetic switching probability. Error bars denote 95% confidence intervals calculated by bootstrapping. Red dashed line represents the average final fitness for purely genetic adaptation.

**Fig S7.**
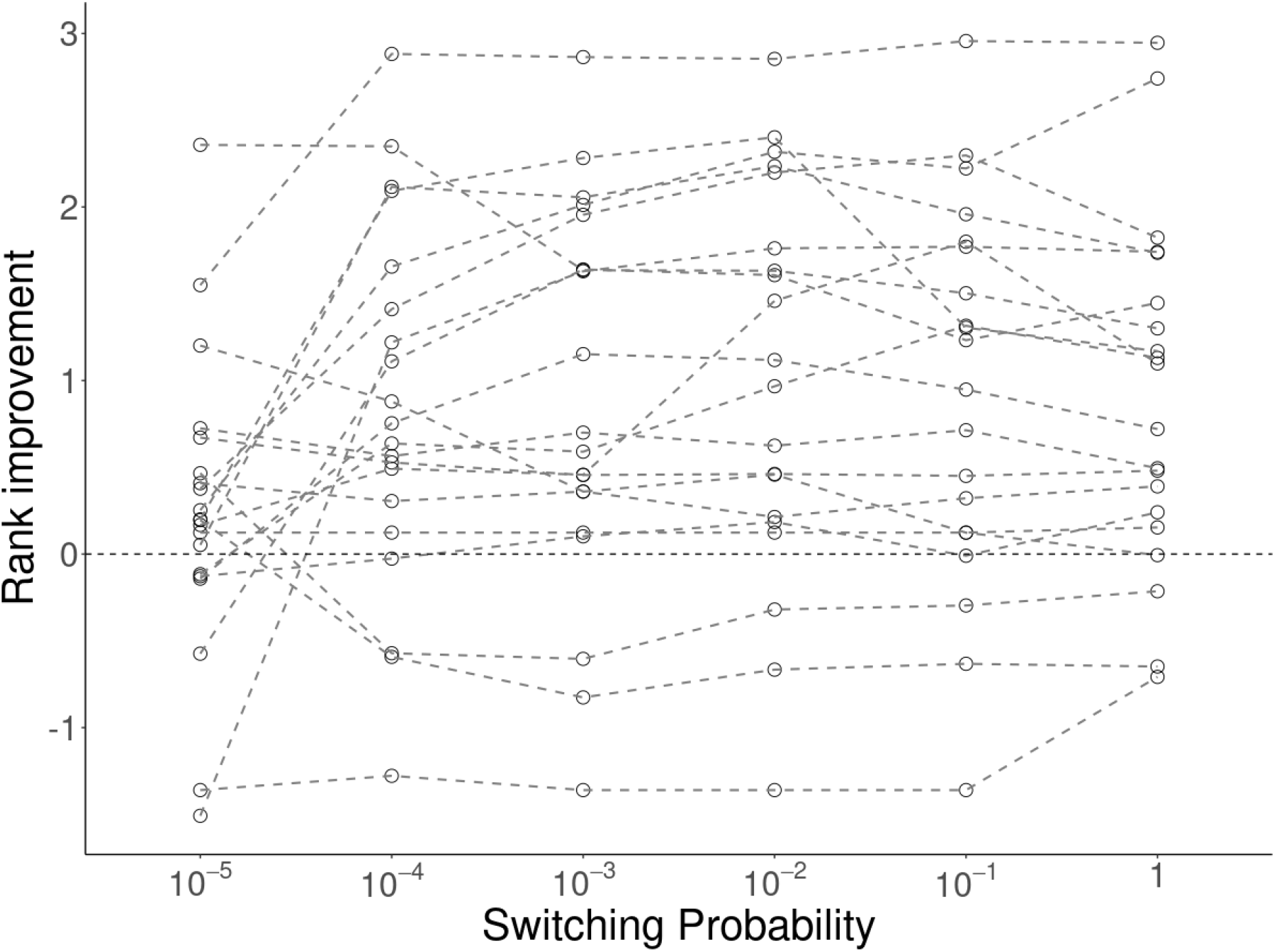
The effect of epigenetics on an individual landscape is independent of epigenetic switching rates. For 20 randomly selected landscapes out of our set of 5-peaked landscapes, we show the effect of epigenetics (rank improvement) as a function of the absolute value of epigenetic switching probabilities (symmetric in both directions here). Dashed lines connect points associated with the same landscape.

